# The Genetic Basis for the Cooperative Bioactivation of Plant Lignans by a Human Gut Bacterial Consortium

**DOI:** 10.1101/357640

**Authors:** Elizabeth N. Bess, Jordan E. Bisanz, Peter Spanogiannopoulos, Qi Yan Ang, Annamarie Bustion, Seiya Kitamura, Diana L. Alba, Dennis W. Wolan, Suneil K. Koliwad, Peter J. Turnbaugh

## Abstract

Plant-derived lignans, consumed daily by most individuals, are inversely associated with breast cancer; however, their bioactivity is only exerted following gut bacterial conversion to enterolignans. Here, we dissect a four-species bacterial consortium sufficient for all four chemical reactions in this pathway. Comparative genomics and heterologous expression experiments identified the first enzyme in the pathway. Transcriptional profiling (RNAseq) independently identified the same gene and linked a single genomic locus to each of the remaining biotransformations. Remarkably, we detected the complete bacterial lignan metabolism pathway in the majority of human gut microbiomes. Together, these results are an important step towards a molecular genetic understanding of the gut bacterial bioactivation of lignans and other plant secondary metabolites to downstream metabolites relevant to human disease.

**One Sentence Summary:** Bess *et al.* provide a first step towards elucidating the molecular genetic basis for the cooperative gut bacterial bioactivation of plant lignans, consumed daily by most individuals, to phytoestrogenic enterolignans.

## Main Text

Plants have been consumed for centuries to prevent and treat disease, yet their medicinal properties can arise not from the food itself but from the metabolites produced by the gut microbiome. The human gut microbiome catalyzes the biotransformation of numerous dietary substrates into bioactive small molecules that can act directly in the gastrointestinal tract or even reach distant tissues following their absorption into general circulation (*1-3*). This fact, together with the tremendous inter-individual variation in gut microbial community structure and function (*4*), highlights the need to design personalized diets for the prevention of human disease (*5*). Yet, this ambitious goal has been difficult to obtain given the major gaps in our knowledge regarding the mechanism and scope of gut microbial metabolism.

Metabolism of lignans to enterolignans, which are one of the most commonly ingested sources of diet-derived phytoestrogens (*6*), occurs exclusively via the action of gut-residing bacteria (*7*). Lignans are ubiquitous in plants and consumed at estimated mean rates of 1-2 milligrams per day, 75% of which are the lignans pinoresinol (PINO) and lariciresinol (LAR) (Fig. 1A) (*8*). However, dietary lignans are poorly bioavailable and have limited bioactivity prior to their activation by human gut bacterial enzymes (*9, 10*). In rats, lignans tend to accumulate in cecal contents, a bacteria-dense region of the gut where significant bacterial metabolism of lignans may occur (*11*). The downstream microbial metabolites, the phytoestrogens enterodiol (END) and enterolactone (ENL), have both structural and functional similarities to the endogenous hormone estrogen and broad impacts for host health and disease (*9, 12*). Most notably, serum ENL levels are inversely associated with breast cancer risk in humans (*13-16*), and ENL dosed to rodent models of breast cancer decreases tumor burden (*17, 18*).

**Fig. 1.**
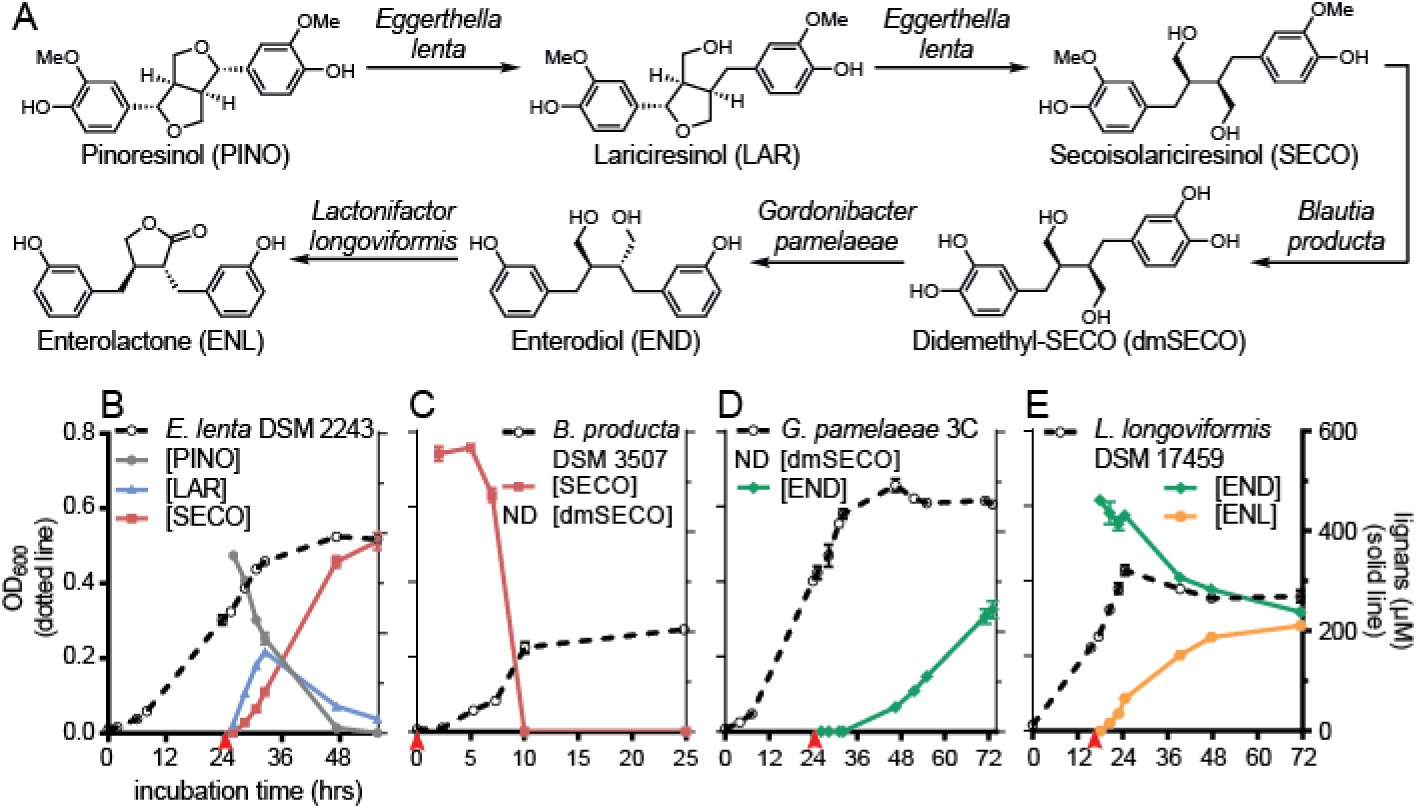
A four-member gut bacterial consortium is capable of converting dietary lignans to phytoestrogenic enterolignans. **(A)** Biotransformations by which gut-residing bacteria can convert the dietarily abundant lignans PINO and LAR into the enterolignan products END and ENL. **(B-E)** Time-course experiments exhibiting the conversion of PINO to ENL and the growth profile of each bacteria responsible for metabolism. Lignan concentrations were measured by HPLC. Due to the absence of commercially available dmSECO standard as well as this metabolite’s instability, dmSECO could not be accurately measured. Culture turbidity, measured as optical density at 600 nm (OD_600_), is depicted in dashed lines. Lignan concentrations, measured by HPLC, are depicted in solid lines. Values are mean±SEM (n=3 biological replicates). Red arrows indicate time at which culture was exposed to lignan.

Despite the discovery of enterolignans in humans nearly 40 years ago, the enzymes responsible for their production are unknown (*19, 20*). Employing a bacterial consortium reported to mediate metabolism of dietary lignans (*7, 21*), here, we used comparative genomics and transcriptional profiling (RNAseq) to link human gut bacterial genetic loci to all four types of chemical transformations involved in the conversion of dietarily abundant PINO and LAR to ENL. The enzymatic activity for the initial two reactions (benzyl ether reductions) was validated by heterologous expression. Together, these results provide new insights into the chemistry made possible by host-associated microbial communities; generalizable approaches to identify genes responsible for the gut microbial metabolism of dietary, therapeutic, or even endogenous small molecules; and an initial step towards the microbiome-based prediction of which individuals would benefit from a lignan-rich diet.

The human gut microbiome metabolizes the dietary lignans PINO and LAR via four distinct types of chemical reactions—benzyl ether reduction, guaiacol demethylation, catechol dehydroxylation, and diol lactonization (Fig. 1A) (*22*)—each carried out by a different bacterial species to yield the final product ENL (*7, 22, 23*). No single gut bacterium has been identified that can catalyze the diverse array of biotransformations constituting the lignan-metabolism pathway. Instead, cooperation appears to be required among multiple species from distinct bacterial phyla to complete the full pathway. Four bacteria, each capable of performing one of the four types of chemistry converting lignans to enterolignans, were cultured under anaerobic conditions with each putative substrate. Consistent with prior studies (*22, 24, 25*), *Eggerthella lenta* DSM 2243 was capable of catalyzing the first two reactions (Fig. 1B), *Blautia producta* DSM 3507 metabolized SECO (Fig. 1C), and *Lactonifactor longoviformis* DSM 17459 converted END to ENL (Fig. 1E). In contrast to prior reports (*7, 22*), END was not detected following the incubation of dmSECO with *E. lenta*; instead, we identified two *Gordonibacter* species, close relatives of *E. lenta*, capable of this reaction (Fig. 1D, Table S1). No significant growth defects were observed upon incubation of each bacterium with its respective substrate (500 µM in methanol).

Although the *E. lenta* type strain (DSM 2243) is capable of metabolizing PINO to SECO via benzyl ether reduction (Fig. 1B), we were curious about whether this phenotype was found in other *E. lenta* strains. We recently reported the curation of 25 strains of the gut Actinobacterium *E. lenta* and closely-related species of the Coriobacteriia class (*26*). Using an HPLC-based assay, this panel of bacteria was screened for the metabolism of PINO to LAR and SECO (Fig. 2). Metabolism varied across strains; 9 strains (36%) did not produce any detectable metabolites (non-metabolizers). Among the 16 metabolizers, the extent of metabolism varied considerably, from 4.8±0.5% to complete conversion of dosed PINO to the reduced products LAR and SECO (Table S2). All of the metabolizing strains were from the *Eggerthella* genus, including 15 *E. lenta* strains and 1 *E. sinensis* strain.

**Fig. 2.**
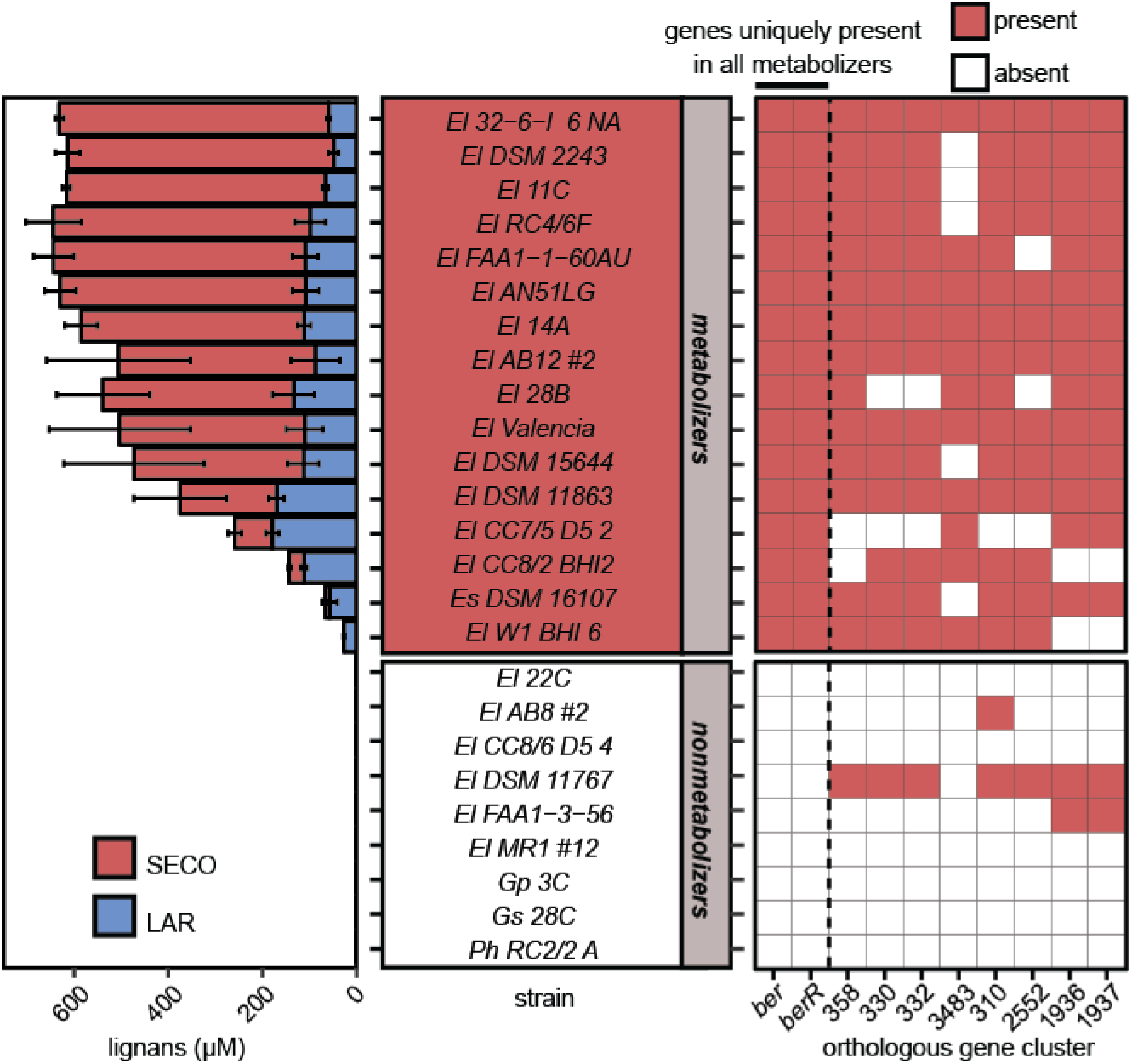
Lignan metabolism varies between Coriobacteriia strains. Evaluation of PINO-metabolizing ability across our collection of strains from the Coriobacteriia class, of which *E. lenta* is a member. Of 25 strains, 16 (all within the *Eggerthella* genus) were found to be metabolizers. Lignans were quantified by HPLC (values are mean±SEM, n=3). Comparative genomic analysis revealed that only two genes, *ber* and *berR*, are present in all PINO-metabolizing strains and absent in all strains that do not metabolize PINO. El: *Eggerthella lenta*; Es: *Eggerthella sinesis*; Gp: *Gordonibacter pamelaeae*; Gs: *Gordonibacter* species; Ph: *Paraeggerthella hongkongensis*.

Phylogenetic analysis revealed that the metabolizing strains were not monophyletic (Fig. S1), de-coupling this phenotypic trait from bacterial evolutionary history and suggesting the repeated gain or loss of the genes responsible. Comparative genomics using our recently developed ElenMatchR tool (*26*) revealed a single genomic locus present in all 16 metabolizing strains and absent in the 9 non-metabolizers (Fig. 2, Table S3). This locus consisted of two genes predicted to encode a putative enzyme and a transcriptional regulator. The former has primary sequence homology to annotated fumarate reductases (Fig. S2) and is referred to herein as benzyl ether reductase (*ber*). The latter, referred to herein as the *ber* regulator (*berR*), has primary sequence homology to annotated transcriptional regulators from the LuxR family. Consistent with its putative regulatory function (*27*), *berR* is transcribed in reverse orientation to *ber*.

As *E. lenta* is presently a genetically intractable organism, we used heterologous expression to confirm that Ber is sufficient to catalyze the benzyl ether reduction of PINO and LAR to SECO. Incubation of *E. coli* Rosetta 2(DE3) cells engineered to express Ber led to 97.2±1.9% conversion of PINO to the products LAR and SECO (Fig. 3A). While there have been reports of genes from sphingomonads (bacteria commonly associated with plant roots) that enable conversion of PINO to LAR and SECO, these genes share no primary sequence homology with *ber* (*28*). To our knowledge, the discovery of *ber* is the first report of a gut bacterial gene that encodes benzyl ether reduction.

**Fig. 3.**
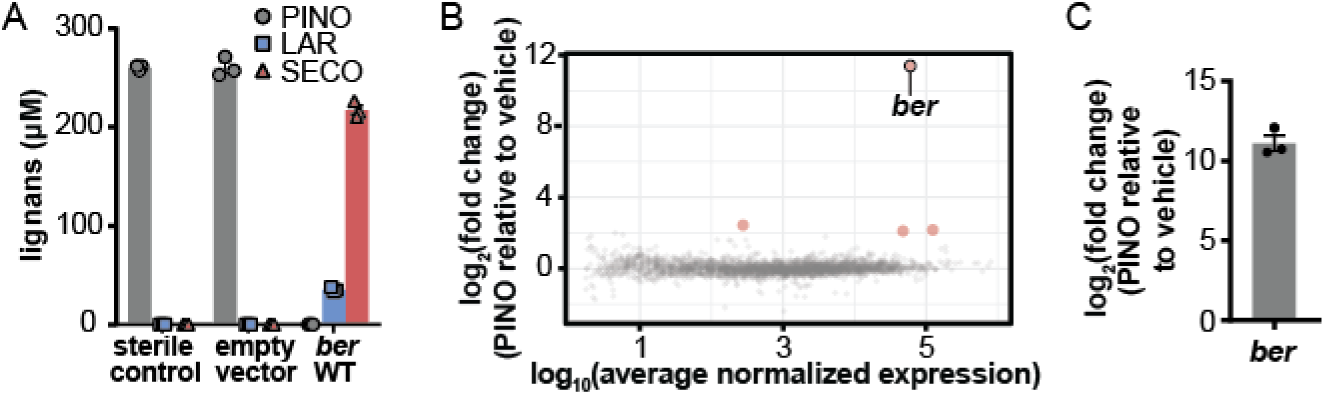
A single enzyme is sufficient to catalyze the first two reactions in the lignan metabolism pathway. **(A)** Incubation of PINO (250 µM) with *E. coli* Rosetta 2(DE3) expressing WT Ber on a pET19btev plasmid. PINO and the products of its benzyl ether reduction, LAR and SECO, were quantified by HPLC (bars are mean±SEM, n=3). **(B)** Plot of RNAseq results obtained upon exposure of *E. lenta* DSM 2243 to PINO relative to vehicle. Points in pink are genes for which was observed a fold-change>|4| and FDR<0.01. **(C)** qRT-PCR confirmation of *ber* up-regulation upon exposure of *E. lenta* DSM 2243 to PINO relative to vehicle (bar is mean±SEM, n=3).

While our comparative genomic approach has been valuable for learning about *E. lenta*, we currently have a limited set of strains for the remaining three bacterial species in the four-membered PINO-to-ENL-metabolizing consortium. As an alternative approach, we turned to transcriptional profiling (RNAseq) to search for genes that are up-regulated in response to each substrate—a strategy that we have successfully employed to implicate genes in the metabolism of pharmaceuticals (*4, 29*).

Incubation of *E. lenta* DSM 2243 with PINO provided an additional validation for this RNAseq-based approach to identify genes that encode xenobiotic-metabolizing activities. The expression of four genes in *E. lenta* DSM 2243 was significantly altered (fold-change>|4|, FDR<0.01) upon exposure to PINO during exponential growth (Fig. 3B; Table S4). The *ber* gene was the most highly up-regulated by a factor of 500 [*ber,* 2670.9-fold; second most up-regulated gene (encoding a hypothetical protein), 5.4-fold]. The induction of *ber* transcription in response to PINO was confirmed by qRT-PCR on an independent set of cultures (Fig. 3C). These results indicate that *ber* transcription is induced upon the cellular detection of Ber’s substrate, PINO, and also suggest that RNAseq may be a general strategy to link genomic loci to specific dietary substrates.

Following the benzyl ether reductions of PINO that produce LAR and then SECO, lignan metabolism proceeds with *Blautia producta* DSM 3507 demethylating the 3-position methoxy groups of SECO, liberating two catechols to form didemethylsecoisolariciresinol (dmSECO) (Fig. 1A, 1C) (*22, 30*). Incubation of SECO with *B. producta* DSM 3507 resulted in the up-regulation of 6 genes and down-regulation of 2 genes (fold-change>|4|, FDR<0.01) (Fig. 4A, Table S5), as determined by RNAseq. The most up-regulated gene (12.6-fold) exhibits sequence homology to known methyltransferases and is referred to herein as guaiacol lignan methyltransferase (*glm*), which putatively mediates demethylation of SECO to yield dmSECO (Fig. 4B, S2). Up-regulation of *glm* was validated by qRT-PCR (Fig. 4B).

**Fig. 4.**
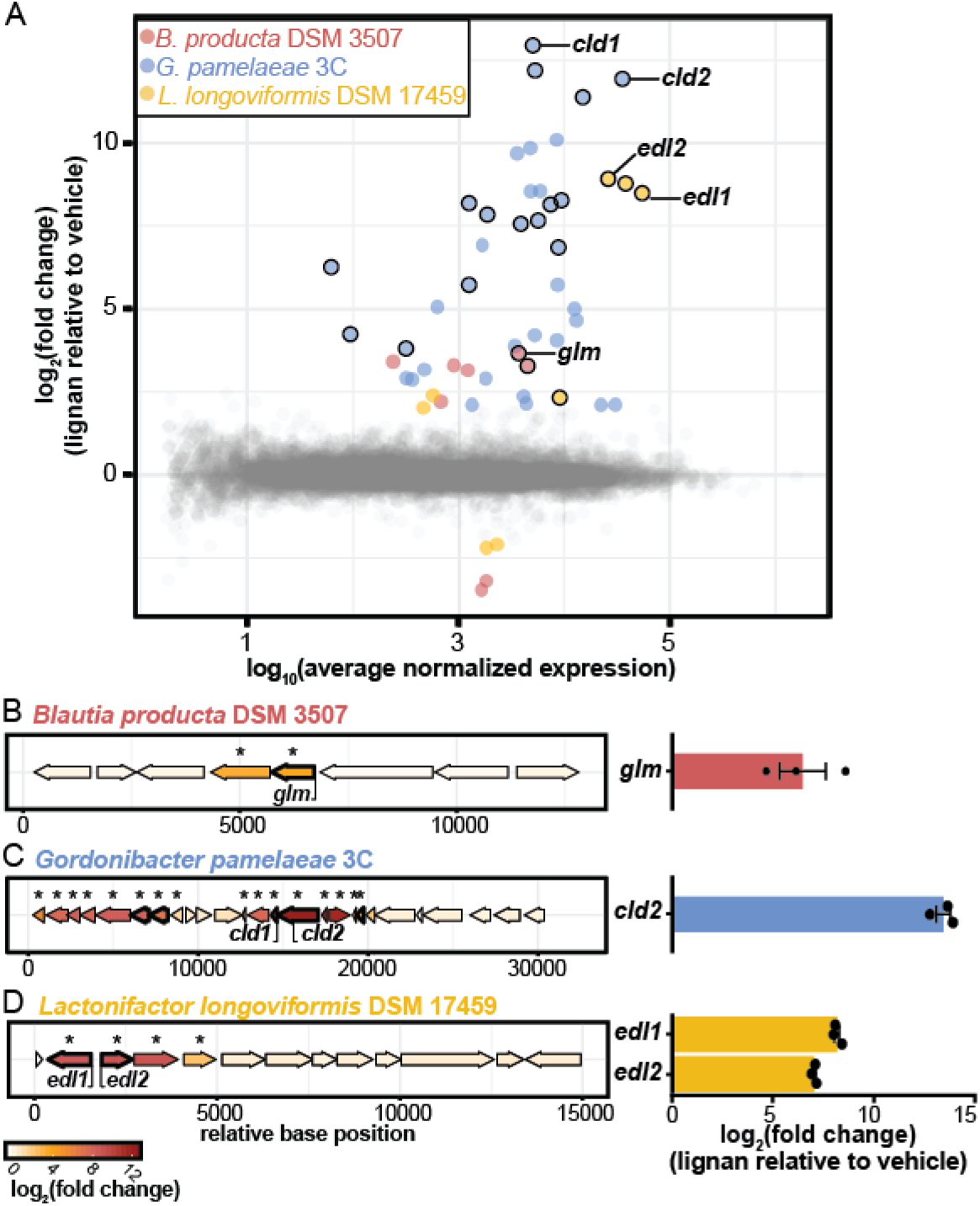
Identification of genomic loci up-regulated in response to each substrate in the lignan metabolism pathway. **(A)** Plot of RNAseq results observed from the exposure of *B. producta* DSM 3507 to SECO, *G. pamelaeae* to dmSECO, and *L. longoviformis* DSM 17459 to END relative to respective vehicle exposures. Colored points have a fold-change>|4| and FDR<0.01. Points encircled with black designate clusters of genes that are represented in the locus diagrams in Fig. 4B-4D. **(B-D)** The most up-regulated genomic loci in *B. producta* DSM 3507, *G. pamelaeae* 3C, and *L. longoviformis* DSM 17459, respectively. Genes that are outlined in bold demonstrate homology to enzymes (Tables S5–S7). The genes implicated in lignan-metabolizing biotransformations are annotated: guaiacol lignan methyltransferase (*glm*), catechol lignan dehydroxylase (*cld*), enterodiol lactonizing enzyme (*edl*). Asterisks denote that the gene is up-regulated in the presence of lignan relative to vehicle with a fold-change>|4| and FDR<0.01. Up-regulation of *glm*, *cld2*, *edl1*, and *edl2* upon exposure to respective lignan substrates relative to vehicle was confirmed through qRT-PCR (bar is mean±SEM, n=3).

The bacterial lignan-metabolism pathway proceeds with *Gordonibacter pamelaeae* 3C dehydroxylating the 4-position hydroxyl group in the catechol moiety of dmSECO, forming END (Fig. 1A, 1D) (*22*). In search of the gene encoding the catechol lignan dehydroxlase (*cld*), RNAseq analysis of *G. pamelaeae* 3C exposed to dmSECO revealed a 19-gene cluster in which 15 genes were significantly up-regulated relative to vehicle (Fig. 4A, Table S6; fold-change>|4|, FDR<0.01). Two proteins encoded in this locus are chemically reasonable candidates for the observed dehydroxylation (Fig. S2). One protein, encoded by the most up-regulated gene (7862.5-fold) in *G. pamelaeae* 3C upon dmSECO exposure, harbors a 4Fe-4S ferredoxin-type binding domain (referred to herein as Cld1). The second protein in this locus is a molybdopterin oxidoreductase (referred to herein as Cld2); *cld2* was up-regulated 3893.8-fold (up-regulation confirmed via qRT-PCR, Fig. 4C). Enzymes bearing molybdenum and iron-sulfur clusters frequently mediate oxygen-transfer reactions (*31*); broadly, this activity is consistent with the catechol dehydroxylation of dmSECO to form END. Both of these genes are only present in the strains of our Coriobacteriia collection that were found to produce END and are absent in all non-metabolizing strains.

Conversion of END to phytoestrogenic ENL by *Lactonifactor longoviformis* DSM 17459, the final step in the lignan-bioactivation pathway (Fig. 1A, 1E) (*25*), was investigated using RNAseq. *L. longoviformis* incubated with END revealed 6 genes that were up-regulated and 2 that were down-regulated in the presence of END relative to vehicle (Fig. 4A, Table S7; fold-change>|4|, FDR<0.01). Among these genes was a single genomic locus of three genes that exhibited transcript levels >300-fold higher in the presence of END (Fig. 4A, 4D); the second most transcriptionally up-regulated gene was only up-regulated 5.3-fold. One of the three dramatically up-regulated genes in this locus is homologous to an MFS transporter. The other two genes in the cluster (up-regulation validated by qRT-PCR, Fig. 4D) demonstrate homology to enzymes from two enzyme super-families: short-chain dehydrogenase/reductases (referred to herein as enterodiol lactonizing enzyme (Edl) 2; *edl2* up-regulation: 470.8-fold) and NADP-dependent oxidoreductases (referred to herein as Edl1; *edl1* up-regulation: 359.6-fold). These genes are plausible sources of the observed oxidative biotransformation.

The prevalence of the bacterial genes herein implicated in the microbiome’s metabolism of dietary PINO to ENL was assessed in a cohort of 68 individuals residing in Northern California (Table S8) (*32*). We developed a novel culture-independent, PCR-based assay for the presence/absence of each gene from community DNA extracted from stool samples (Supplementary Methods). The individual genes varied in their prevalence: *ber*, 93%; *glm*, 87%; *cld2*, 81%; *edl1*, 96%; and *edl2*, 99% (Fig. 5). We did not detect any significant correlations between age, ethnicity, or body mass index and the presence of these genes. Remarkably, we detected the complete bacterial pathway for lignan metabolism in the majority (67.6%) of individuals, suggesting that this is a common mechanism by which the gut microbiome produces phytoestrogenic enterolignans.

**Fig. 5.**
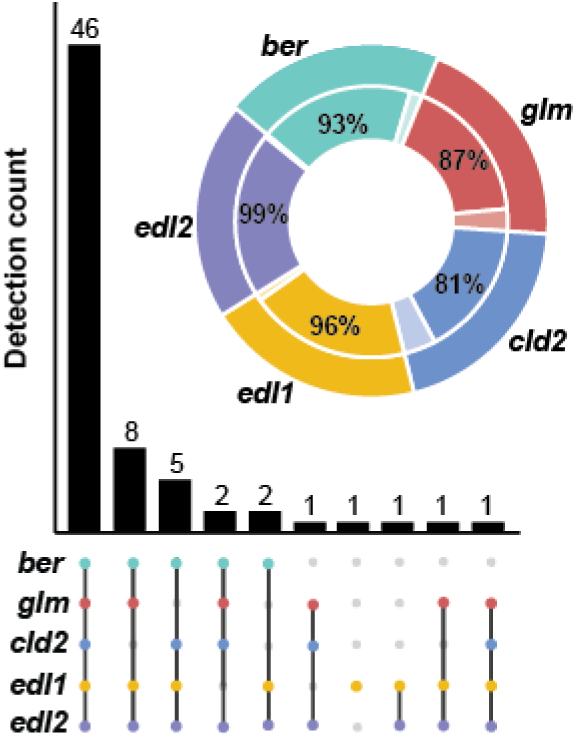
The bacterial lignan metabolism pathway is detectable in the majority of human gut microbiomes. The pie chart displays the prevalence of *ber*, *glm*, *cld2*, *edl1*, and *edl2*, as detected by PCR, in the stool samples of 68 adults living in Northern California (Table S8). The “UpSet” plot (*34*) represents within-sample patterns in gene co-occurrence and the number of samples in which that pattern is detected (detection count). 46/68 individuals (67.6%) are positive for all 5 genes.

While bacterial species that participate in the multi-step, multi-organism metabolism of dietary lignans had previously been reported (*22*), our results demonstrate that strain-level genetic variation within the *E. lenta* population precludes prediction of lignan-metabolizing ability from a species-level depiction of the microbiome. This strain-level variation can be leveraged through the use of comparative genomics to identify genetic determinants of lignan metabolism, as reported previously for antibiotic resistance (*26*) and drug metabolism (*33*). Our finding that *ber* was dramatically, uniquely up-regulated upon exposure to its PINO substrate inspired the application of transcriptional-profiling (RNAseq) to probe the other three steps in the bacterial production of enterolignans. Together, these results emphasize the utility of RNAseq as a generalizable strategy for linking genomic loci to metabolic activities (*4, 29*), even in genetically intractable microorganisms. We anticipate that these two approaches could aid in efforts to gain a more systematic understanding of the full scope of chemical reactions that are enacted by the human microbiome.

Our results enabled the design of a simple culture-independent test for the capacity of the gut microbiome to produce enterolignans, and thus impact host cellular pathways, on a lignan-rich diet. Continued refinement of this method and its functional validation would enable detecting and filling of gaps in key bacterial metabolic pathways through the selective introduction of specific strains to the gut microbiome. More broadly, these results set the stage for an in-depth analysis of the biochemistry of each of these enzymes (Fig. S3), which together act on conserved chemical motifs that are found in numerous dietary, pharmaceutical, and endogenous small molecules.

## Acknowledgments

Special thanks to Emily Balskus, Andrew Patterson, and Katie Pollard for Acknowledgments: Special thanks to Emily Balskus, Andrew Patterson, and Katie Pollard for Graduate Program) and Emily Waligurski (UCSF Summer Research Training Program) for technical assistance and to Separation Research Ltd. (Turku, Finland) for the donation of chemicals.

## Author contributions

P.J.T.; Methodology, E.N.B. and J.E.B.; Software, J.E.B.; Investigation, E.N.B., J.E.B., P.S., A.B., and Q.A.; Resources, S.K., D.L.A., D.W.W., S.K.K. Writing – Original Draft, E.N.B. Writing – Review & Editing, E.N.B. and P.J.T. Supervision, E.N.B. and P.J.T. Funding Acquisition, E.N.B. and P.J.T.

## Data and materials availability

All data and code is available in the main text or supplemental materials, with the exception of RNA-sequencing data, which is available in NCBI’s Sequence Read Archive, accession number SRP140684.

## Supplementary Materials

Materials and Methods 5

Figures S1-S3

Tables S1-S11

CodeS1.html

## References and Notes

1. P. Spanogiannopoulos, E. N. Bess, R. N. Carmody, P. J. Turnbaugh, The microbial pharmacists within us: a metagenomic view of xenobiotic metabolism. Nat. Rev. Microbiol. 14, 273–287 (2016).

2. R. N. Carmody, P. J. Turnbaugh, Host-microbial interactions in the metabolism of therapeutic and diet-derived xenobiotics. J. Clin. Invest. 124, 4173–4181 (2014).

3. N. Koppel, V. Maini Rekdal, E. P. Balskus, Chemical transformation of xenobiotics by the human gut microbiota. Science 356, (2017).

4. C. F. Maurice, H. J. Haiser, P. J. Turnbaugh, Xenobiotics shape the physiology and gene expression of the active human gut microbiome. Cell 152, 39–50 (2013).

5. R. Jumpertz von Schwartzenberg, P. J. Turnbaugh, Siri, What Should I Eat? Cell 163, 1051–1052 (2015).

6. L. M. Valsta et al., Phyto-oestrogen database of foods and average intake in Finland. Br. J. Nutr. 89, S31–S38 (2011).

7. A. Woting, T. Clavel, G. Loh, M. Blaut, Bacterial transformation of dietary lignans in gnotobiotic rats. FEMS Microbiol. Ecol. 72, 507–514 (2010).

8. I. Tetens et al., Dietary intake and main sources of plant lignans in five European countries. Food Nutr. Res. 57, (2013).

9. T. Clavel, J. Doré, M. Blaut, Bioavailability of lignans in human subjects. Nutr. Res. Rev. 19, 187–196 (2006).

10. T. Clavel, J. O. Mapesa, in Natural Products: Phytochemistry, Botany and Metabolism of Alkaloids, Phenolics and Terpenes, K. G. Ramawat, J.-M. Mérillon, Eds. (Springer Berlin Heidelberg, Berlin, Heidelberg, 2013), pp. 2433–2463.

11. S. E. Rickard, L. U. Thompson, Chronic exposure to secoisolariciresinol diglycoside alters lignan disposition in rats. J. Nutr. 128, 615–623 (1998).

12. H. Adlercreutz, Lignans and human health. Crit. Rev. Clin. Lab. Sci. 44, 483–525 (2007).

13. A. K. Zaineddin et al., Serum enterolactone and postmenopausal breast cancer risk by estrogen, progesterone and herceptin 2 receptor status. Int. J. Cancer 130, 1401–1410 (2012).

14. A. Olsen et al., Plasma enterolactone and breast cancer incidence by estrogen receptor status. Cancer Epidemiol. Biomarkers Prev. 13, 2084–2089 (2004).

15. E. Sonestedt et al., Enterolactone is differently associated with estrogen receptor β-negative and -positive breast cancer in a Swedish nested case-control study. Cancer Epidemiol. Biomarkers Prev. 17, 3241–3251 (2008).

16. P. Seibold et al., Enterolactone concentrations and prognosis after postmenopausal breast cancer: Assessment of effect modification and meta-analysis. Int. J. Cancer 135, 923–933 (2014).

17. H. B. Mabrok et al., Lignan transformation by gut bacteria lowers tumor burden in a gnotobiotic rat model of breast cancer. Carcinogenesis 33, 203–208 (2012).

18. N. M. Saarinen et al., Enterolactone inhibits the growth of 7,12-dimethylbenz(a) anthracene-induced mammary carcinomas in the rat. Mol. Cancer Ther. 1, 869–876 (2002).

19. S. R. Stitch et al., Excretion, isolation and structure of a new phenolic constituent of female urine. Nature 287, 738–740 (1980).

20. K. D. R. Setchell et al., Lignans in man and in animal species. Nature 287, 740–742 (1980).

21. T. Clavel et al., Intestinal bacterial communities that produce active estrogen-like compounds enterodiol and enterolactone in humans. Appl. Environ. Microbiol. 71, 6077–6085 (2005).

22. T. Clavel, D. Borrmann, A. Braune, J. Doré, M. Blaut, Occurrence and activity of human intestinal bacteria involved in the conversion of dietary lignans. Anaerobe 12, 140–147 (2006).

23. A. K. Bolvig et al., Effect of antibiotics and diet on enterolactone concentration and metabolome studied by targeted and nontargeted LC-MS metabolomics. J. Proteome Res. 16, 2135–2150 (2017).

24. L. Q. Wang, M. R. Meselhy, Y. Li, G. W. Qin, M. Hattori, Human intestinal bacteria capable of transforming secoisolariciresinol diglucoside to mammalian lignans, enterodiol and enterolactone. Chem. Pharm. Bull. 48, 1606–1610 (2000).

25. T. Clavel, G. Henderson, W. Engst, J. Doré, M. Blaut, Phylogeny of human intestinal bacteria that activate the dietary lignan secoisolariciresinol diglucoside. FEMS Microbiol. Ecol. 55, 471–478 (2006).

26. J. E. Bisanz et al., Illuminating the microbiome’s dark matter: a functional genomic toolkit for the study of human gut Actinobacteria. *bioRxiv*, (2018).

27. R. M. Davis, R. Y. Muller, K. A. Haynes, Can the natural diversity of quorum-sensing advance synthetic biology? Front. Bioeng. Biotechnol. 3, (2015).

28. Y. Fukuhara et al., Discovery of pinoresinol reductase genes in sphingomonads. Enzyme Microb. Technol. 52, 38–43 (2013).

29. H. J. Haiser et al., Predicting and manipulating cardiac drug inactivation by the human gut bacterium Eggerthella lenta. Science 341, 295–298 (2013).

30. L.-Q. Wang, M. R. Meselhy, Y. Li, G.-W. Qin, M. Hattori, Human intestinal bacteria capable of transforming secoisolariciresinol diglucoside to mammalian lignans, enterodiol and enterolactone. Chem. Pharm. Bull. 48, 1606–1610 (2000).

31. R. Hille, The molybdenum oxotransferases and related enzymes. Dalton Trans. 42, 3029–3042 (2013).

32. D. L. Alba et al., Subcutaneous fat fibrosis links obesity to insulin resistance in Chinese-Americans. J. Clin. Endocrinol. Metab., jc.2017–02301 (2018).

33. N. Koppel, J. E. Bisanz, M.-E. Pandelia, P. J. Turnbaugh, E. P. Balskus, Discovery and characterization of a prevalent human gut bacterial enzyme sufficient for the inactivation of a family of plant toxins. eLife 7, e33953 (2018).

34. J. R. Conway, A. Lex, N. Gehlenborg, UpSetR: an R package for the visualization of intersecting sets and their properties. Bioinformatics 33, 2938–2940 (2017).

